# Image-guided deployment and monitoring of a novel tungsten nanoparticle–infused radiopaque absorbable inferior vena cava filter in pigs

**DOI:** 10.1101/2023.02.06.527049

**Authors:** Erin Marie San Valentin, Jossana A. Damasco, Marvin Bernardino, Karem A. Court, Biana Godin, Gino Martin Canlas, Adam Melancon, Gouthami Chintalapani, Megan C. Jacobsen, William Norton, Rick R. Layman, Natalie Fowlkes, Stephen R. Chen, Steven Y. Huang, Marites P. Melancon

## Abstract

The use of absorbable inferior vena cava filters (IVCFs) constructed with poly-p-dioxanone (PPDO) eliminates risks and complications associated with the use of retrievable metallic filters. Radiopacity of radiolucent PPDO IVCFs can be improved with the incorporation of nanoparticles (NPs) made of high-atomic number materials such as gold and bismuth. In this study, we focused on incorporating tungsten NPs (WNPs), along with polyhydroxybutyrate (PHB), polycaprolactone (PCL), and polyvinylpyrrolidone (PVP) polymers to increase the surface adsorption of the WNPs. We compared the imaging properties of WNPs with single-polymer PHB (W-P) and WNPs with polymer blends consisting of PHB, PCL, and PVP (W-PB). Our *in vitro* analyses using PPDO sutures showed enhanced radiopacity with either W-P or W-PB coating, without compromising the inherent physico-mechanical properties of the PPDO sutures. We observed a more sustained release of WNPs from W-PB-coated sutures than W-P-coated sutures. We successfully deployed W-P- and W-PB-coated IVCFs into the inferior vena cava of pig models, with monitoring by fluoroscopy. At the time of deployment, W-PB-coated IVCFs showed a 2-fold increase in radiopacity compared to W-P-coated IVCFs. Longitudinal monitoring of *in vivo* IVCFs over a 12-week period showed a drastic decrease in radiopacity at week 3 for both filters. Results of this study highlight the utility of NPs and polymers for enhancing radiopacity of medical devices; however, different methods of incorporating NPs and polymers can still be explored to improve the efficacy, safety, and quality of absorbable IVCFs.

## Introduction

Deep vein thrombosis medical condition involving the formation of blood clots in a deep vein, and may migrate to the lungs, where it is then referred to as pulmonary embolism (PE). Though highly preventable, this medical condition is often underdiagnosed and serious, which may lead to disability and death. At least 900,000 people are estimated to have DVT or PE in the United States alone.^1^ Typically, anticoagulants and fibrinolytics are prescribed to reduce or dissolve existing clots. However, for patients with contraindications to anticoagulation, placement of inferior vena cava filters (IVCFs) is recommended. These devices are placed in the inferior vena cava to trap clots and keep them from passing into the lungs. IVCFs can be permanent or temporary, with an intended dwell time of 7 to 35 days.^2, 3^ Non-retrieval of temporary IVCFs poses patient safety concerns, including risks of IVCF migration, embolization, and thrombosis.^4^

Absorbable IVCFs constructed with poly-p-dioxanone (PPDO) eliminate the need for filter retrieval.^5, 6^ The favorable degradation profile of PPDO IVCFs allows for desired mechanical strength and integrity during the intended dwell time, followed by absorption after the intended use. Using absorbable IVCFs could enhance safety and decrease medical cost without compromising the efficacy of traditional IVCFs. However, PPDO IVCFs are radiolucent, making fluoroscopy-guided filter deployment and long-term monitoring difficult. To improve visualization, the addition of radio-enhancing nanoparticles (NPs) such as gold (Au) NPs^3, 7^ has been explored. Our group successfully demonstrated the safety and efficacy of AuNP-coated IVCFs in pigs.^3^ However, due to the high cost of Au, alternative NPs needed to be explored.

Our group then investigated the material differentiation of potential contrast materials such as Au, bismuth (Bi), tungsten (W), ytterbium, gadolinium, iron, tantalum, zirconium, and barium using computed tomography (CT)–based quantification methods.^8^ High-Z (i.e., a high atomic number, >73) NPs are ideal, because they can be differentiated from iodinated agents, which is valuable because iodine contrast media are used during IVCF deployment to assess and monitor intravenous blood clots caught by the filter. Among the potential materials, Au, Bi and W consistently showed properties of a good contrast material for CT imaging in our preliminary tests. These data led to our group’s creation of BiNP-coated IVCFs^9^ which also established that, similar to AuNPs, BiNPs can be used as a radioenhancer for PPDO. To increase the adsorption of BiNPs on the surface of the PPDO absorbable filter, we used polyhydroxybutyrate (PHB). Results showed that PHB reinforcement significantly improved image-guided filter deployment and long-term radiopacity compared to PPDO IVCFs and BiNP-PPDO IVCFs alone. Results of this study suggested that the addition of polymers to NP coatings augments the adsorption of NPs.

Based on those results, we tested the efficacy of tungsten (W) NPs with polymer blends in enhancing the radiopacity of inherently radiolucent PPDO filters. Similar to Bi and Au, W is also a high-Z element (Z=74) with K-edge = 69.5 keV. More importantly, its CT intensity for different X-ray spectra remained steady throughout all energies tested – a trend similar to that observed with Au and Bi. In this study, we investigated the feasibility of W with a single polymer and polymer blends as an alternative NP coating for absorbable PPDO IVCFs in pigs over a 12-week period. We evaluated the effects of WNPs and polymers on the mechanical integrity and clot-trapping efficacy of PPDO IVCFs.

## Experimental section

### Materials

WebMax PPDO sutures (violet monofilament, D451; size 2-0) were obtained from Patterson Veterinary (Greeley, CO). Tungstic acid (99+%) was purchased from Thermo Fisher Scientific (Waltham, MA). Dichloromethane (DCM, ACS reagent grade, ≥99.5%), oleylamine (technical grade, 70%), oleic acid (technical grade, 90%), and 1-octadecene (technical grade, 90%) were obtained from Sigma-Aldrich (St. Louis, MO). Polyvinylpyrrolidone (PVP, average MW 40,000), polyhydroxybutyrate (PHB, granules), and polycaprolactone (PCL) were purchased from Sigma-Aldrich. All chemicals were used without further purification unless otherwise noted.

### Synthesis and characterization of WNPs

WNPs were synthesized by a one-pot thermal decomposition of 1.25 mmol tungstic acid with 30 mL oleic acid, 30 mL oleylamine, and 50 mL 1-octadecene as stabilizers in a 500-mL three-necked round-bottom flask. The mixture was degassed with continuous argon flow at 70°C for 30 min, before heating up to 330°C for another 30 min under argon gas protection. The mixture was cooled to room temperature before collection of blue precipitate consisting of WNPs. Collected NPs were washed with ethanol and stored and dispersed in chloroform prior to use. Collected WNPs were visualized and characterized by transmission electron microscopy (TEM) using a JEM 1010 microscope (JEOL USA Inc., Peabody, MA). The size and distribution of WNPs were measured using ImageJ software (National Institutes of Health, Bethesda, MD).

### Coating and characterization of WNP-PPDO sutures

Similar to our previous studies^3, 9^, we characterized NP-coated or uncoated sutures. WebMax PPDO monofilament sutures were cut into 30-cm lengths and soaked in DCM containing WNPs at a concentration of 0.1 mmol in 1 mL of DCM. WNPs with single-polymer PHB (W-P) and WNPs with polymer blends consisting of PHB, PCL, and PVP (W-PB) were prepared. We characterized W-P- and W-PB-coated sutures using micro-computed tomography (microCT) using a SkyScan 1276 system (Bruker, Billerica, MA); a field-emission scanning electron microscope (FE-SEM) (Nova NanoSEM 230, FEI Company, Hillsboro, OR); and energy-dispersive X-ray spectroscopy (EDX) (EDAX Element EDS, AMETEK Inc., Berwyn, PA) analyzed using TEAM WDS Analysis System software (version V4.5.1-RC13). Tensile strength of coated and uncoated sutures was also measured using an eXpert 7601 Tension Testing System (ADMET Inc., Norwood, MA).

To monitor WNP release from the polymers *in vitro*, control PPDO, W-P-coated, and W-PB-coated sutures were soaked in phosphate buffered saline (PBS) at pH 7.4 and 37°C. The PBS solution was collected and changed at weeks 0, 3, 5, 6, 8, 10, and 12. Tungsten dissolution was performed as described in literature^10^; solutions were mixed with 1M hydrogen peroxide (pH=3) and incubated at 60°C for 30 min for optimum dissolution. An aliquot of 1 mL was obtained and diluted with 2% nitric acid before running for elemental analysis using an inductively coupled plasma optical emission spectrometer (ICP-OES, Varian 720-ES, Agilent, Santa Clara, CA). For each run, we used yttrium as the internal control at 371.03 nm, while W was detected at 207.91, 220.45, and 209.48 nm. Intensity values were plotted against the standard curve in order to obtain the concentration of WNP released in solution.

### Infusion of WNPs onto PPDO IVCFs

PPDO IVCFs were obtained from Adient Medical (Pearland, TX). As previously characterized^5^, the filters measured 47 mm (length) x 20 mm (diameter) and included a stent portion and a conical portion (the capture basket). The stent end was fitted with a radiopaque tip composed of biodegradable polymer containing platinum-iridium. To serve as anchors and radiopaque markers, two 2-mm stainless steel barbs were crimped around the circumference of the stent.

Similar to the infusion of PPDO sutures, IVCFs were coated with W-P or W-PB using a wet-dipping technique. Radiopacity was evaluated using microCT and X-ray scanners. Prior to deployment, IVCFs were subjected to ethylene oxide sterilization.

### Filter deployment and administration of autologous thrombus in pig models

Five adult pigs (35-45 kg at the beginning of the study) were obtained from MD Anderson’s Keeling Center for Comparative Medicine and Research (Bastrop, TX). Protocols used for the study were approved by the MD Anderson Institutional Animal Care and Use Committee. Animals were maintained in facilities approved by AAALAC (Association for Assessment and Accreditation of Laboratory Animal Care) International and in accordance with current U.S. Department of Agriculture, Department of Health and Human Services, and National Institutes of Health regulations and standards.

Following a previously described procedure of filter implantation,^5, 6^ pigs were sedated with tiletamine/zolazepam(4.4 mg/kg) and xylazine (2 mg/kg). Isoflurane (5%), buprenorphine (0.04 mg/kg), and ketoprofen (3 mg/kg) was used to anesthetize the pigs, and maintained with isoflurane (1.5-3%) and oxygen (1.5-2 L/min). Heparin (50 U/kg) was administered prior to catheterization. An Angio/CT combination suite (Miyabi, Siemens, Erlangen, Germany) with a single-plane C-arm digital subtraction angiography (DSA) unit and multidetector CT unit was used for all animal procedures. Filter deployment was initiated by accessing the right femoral vein to the right jugular vein, and an over-the-wire pusher rod was used to advance the filter in an introducer. The sheath was pulled back to expose the filter, and a balloon (Coda K032869, Cook Medical) was advanced and inflated for 5-10 s over the wire. Upon removing the balloon, a pigtail catheter was reintroduced into the right iliac vein, and an inferior venacavogram was performed.

Autologous thrombus was administered immediately upon filter deployment. A post-deployment venacavogram and CT scans of the chest and abdomen/pelvis were taken to evaluate the extent of deployed thrombus and to monitor any evidence of PE.

### *In vivo* imaging and data processing

*In vivo* imaging and data processing was performed as described in our previous publication on BiNP IVCFs ^9^. Briefly, CT images were taken at weeks 0, 3, 5, 6, 8, 10, and 12 using a Siemens SOMATOM Definition Edge scanner (Siemens Healthineers, Forchheim, Germany) (120 kV, 80 mA, slice thicknesses of 1.5 and 3 mm) with or without intravenous iodixanol contrast medium (320 mg/mL; GE Healthcare, Little Chalfont, UK) and evaluated for IVCF radiopacity, clot-trapping ability, IVC thrombosis, caval narrowing, and PE.

CT images were reconstructed through syngo.via software (Siemens Medical Solutions, Malvern, PA) and analyzed using the built-in MM Oncology protocol with a 3-mm slice thickness, then exported to ImageJ software. Regions of interest encompassing the IVC and the implanted IVCFs were created using ImageJ, and the maximum grayscale values within these regions were measured. The grayscale values were taken throughout all slices containing the IVCFs and were averaged to calculate the HU values of the IVCFs. The areas with metal markers were not included in the measurements to avoid a false-positive signal.

### Serial hematologic tests

All pigs underwent routine hematologic tests with blood collected at the time of imaging for analyses of serum chemistry, complete blood counts, coagulation profiles, liver function enzymes, and arterial blood gasses. Blood samples were analyzed at the MD Anderson Keeling Center for Comparative Medicine and Research.

### Necropsy, biodistribution, and histological analysis

Pigs were euthanized by exsanguination under deep anesthesia at 12 weeks after filter deployment. All animals underwent necropsy, and the IVC, lungs, heart, liver, kidneys, spleen, and small and large intestines were collected. Harvested tissues (approximately 1 g each) were weighed and placed in 50-mL conical tubes until further use. Tissues were homogenized using a Polytron PT 2500 E (Kinematica Inc., Bohemia, NY) before digestion with hydrogen peroxide using the same reaction described above. After digestion, an aliquot of 1 mL was diluted to 10 mL with 2% nitric acid and passed through a 0.2-µm filter. W content was quantified in triplicate using an ICP-OES.

Histological evaluation was done by dividing the IVC into two portions; the upper portion was evaluated for incorporation within the vessel wall, integrity, and the degree of filter resorption, while the lower portion was assessed for histological changes with hematoxylin and eosin (H&E) staining. Polarized light microscopy was used to confirm the presence of PPDO suture material in the IVC.

### Statistical analysis

GraphPad Prism version 9.0.0 was used to analyze the data presented in this study. Data were presented as means ± standard deviations and were analyzed using one-way analysis of variance and/or two-tailed Student *t*-test, following the assumption of a normal distribution. For correction of multiple, comparisons, we used the Holm-Sidak method. A p-value less than 0.05 was used to determine significant statistical differences.

## Results and Discussion

### Synthesis and characterization of WNPs

Precise image-guided deployment of IVCFs increases procedural success rate and safe implantation while mitigating the risks of procedure-related complications. Through enhancing the radiopacity of radiolucent PPDO IVCFs, recognition of malposition, filter tilt, and wall apposition would be faster and more efficient without increasing fluoroscopy time. Moreover, improved radiopacity also allows for long-term monitoring of IVCF integrity *in vivo*. While our group previously evaluated the performance of AuNPs and BiNPs with PHB-coated IVCFs, we aim to further improve longitudinal radiopacity by selecting alternative NPs as potential contrast agents in combination with different polymer blends. Our group previously characterized the material decomposition of potential contrast agents including Bi, Au, and W.^8^ W demonstrated a sustained attenuation throughout different X-ray energies and has higher absorption coefficient (4.438 cm^2^/kg at 100 keV) compared to iodine (1.94 cm^2^/kg at 100 keV),^11^ making it a suitable candidate for a novel coating for radiolucent PPDO IVCFs. Although nanostructured tungsten oxide has already been developed and used for electrochromic devices,^12^ photocatalysts,^13^ and field emission displays,^14^ its current biomedical application has been limited. Zhou *et al*.^11^ demonstrated the use of tungstic oxide nanorods with polyethylene glycol (PEG) for simultaneous CT imaging and near infrared photothermal therapy of tumors *in vivo*. The antimicrobial activities of tungsten nanodots^15^ and nanocomposite fibers^16^ have also been explored recently.

In our study, one-pot thermal decomposition of tungstic acid was used to synthesize WNPs. This method allows for single-step large-scale production of WNPs while also controlling the size and morphology of synthesized NPs. The successfully synthesized crystal-like, plate-shaped WNPs were visualized by TEM, showing WNPs of 16 ± 3.73 nm in length by 6.48 ± 1.96 nm in width (Fig 1).

**Fig 1.**
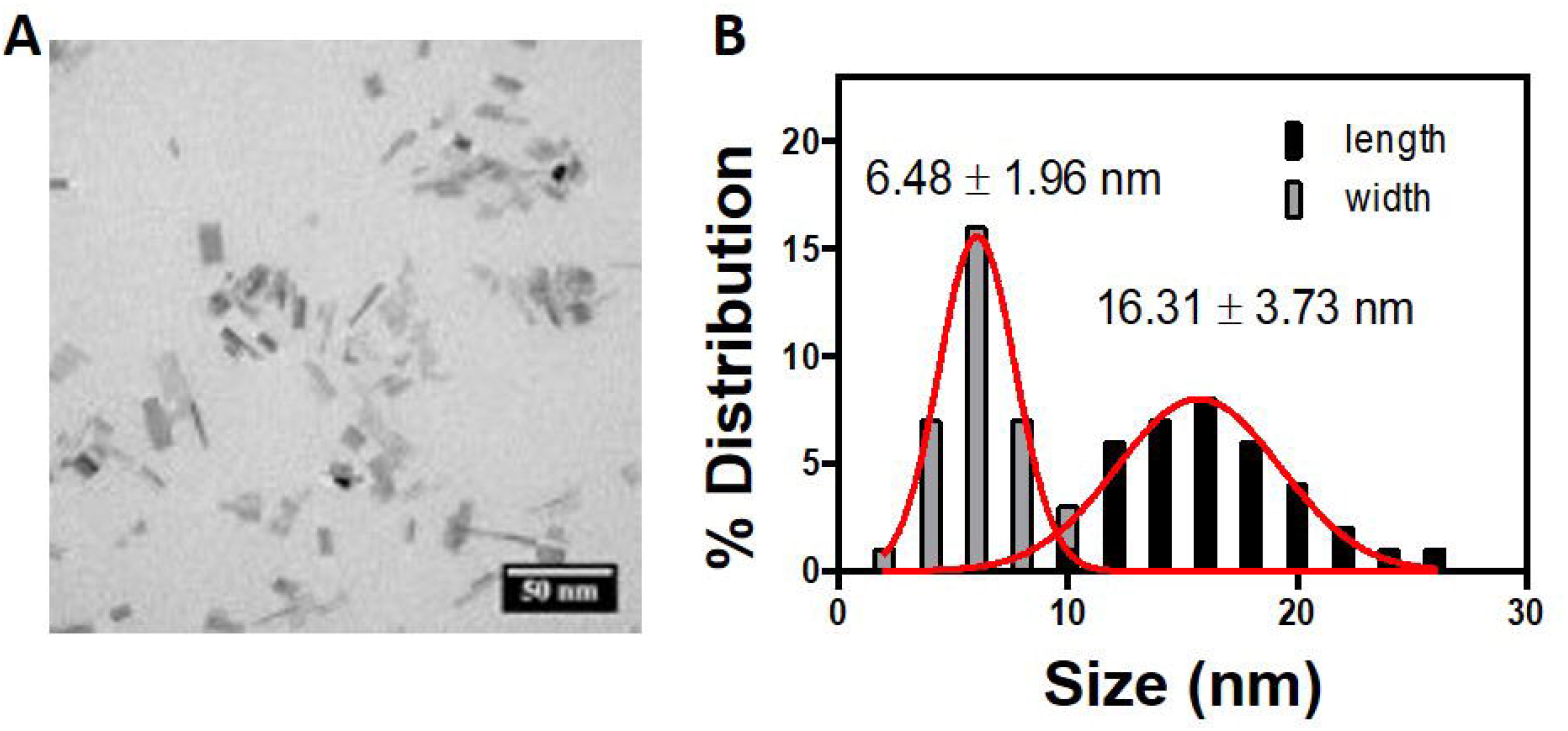
Synthesis and characterization of WNPs. (A) TEM image of synthesized WNPs. (B) Size distribution of WNPs from TEM image.

### Characterization of W-P- and W-PB-infused PPDO sutures

We previously established that a BiNP coating reinforced with PHB increases surface roughness, resulting in increased radiopacity compared to control PPDO and BiNP coating alone.^9^ PHB is a biodegradable polymer and is considered safe for use in medical devices because its monomer, 3-hydroxybutyric acid, is a natural component of human blood.^17^ It has been used in U.S. FDA-approved surgical sutures. Here, we tested its efficacy in enhancing the infusion of WNPs (W-P). Similar to our previous studies, a wet-dipping method was done to coat monofilament PPDO sutures with W-P solutions. However, instead of alternately dipping and drying between NP and polymer solutions, polymer solutions in DCM were mixed with the WNPs. The mixture allowed for a more uniform coating of polymers and NPs onto the PPDO sutures. Our results show that W-P (99.78 ± 13.58 Hounsfield units, HU) enhanced radiopacity of control PPDO (33.72 ± 8.15 HU) by only 33% (Table 1). Therefore, we sought to further enhance the infusion of WNPs within the PPDO sutures. The degradable polymers PCL and PVP were tested to further enhance the radiopacity of PPDO IVCFs. Both polymers have been established to be biodegradable, stable, and biocompatible.^18, 19^ While PCL is commonly used in tissue engineering applications and surgical implants,^20^ PVP has been mainly used as a hydrogel coating in biomedical applications such as contact lenses^21^ and wound dressings.^19^ We then proceeded with coating the PPDO sutures with W-PB solutions. W-PB-coated sutures (0.401 ± 0.000 mm) were thicker than W-P (0.384 ± 0.003 mm) alone, owing to the additional layers of PHB, PVP, and PCL. To confirm the presence of WNPs in W-P- and W-PB-coated sutures, we used SEM with EDX. The clear peaks seen at 1.77 keV (Fig 2) confirmed the presence of W in the W-P- and W-PB-coated sutures. These W peaks can be seen between those for carbon (0.277 keV), oxygen (0.523 keV), and platinum (2.05 keV), which was used for sputter coating of the sutures.

**Table 1.** Physicochemical properties of synthesized WNPs.

|  | % W (wt/wt) | T<sub>m</sub><br>(°C) | Suture Thickness<br>(mm) | Maximum Stress<br>(mPa) | Radiopacity (HU) |
| --- | --- | --- | --- | --- | --- |
| Control PPDO | - | 105.93 ± 0.34 | 0.385 ± 0.001 | 32.56 ± 5.63 | 33.72 ± 8.15 |
| W-P | 25 | 105.87 ± 3.10 | 0.384 ± 0.003 | 37.04 ± 4.56 | 99.78 ± 13.58 |
| W-PB | 15 | 107.94 ± 0.61 | 0.401 ± 0.000 | 31.70 ± 4.12 | 2132.19 ± 323.44 |

**Fig 2.**
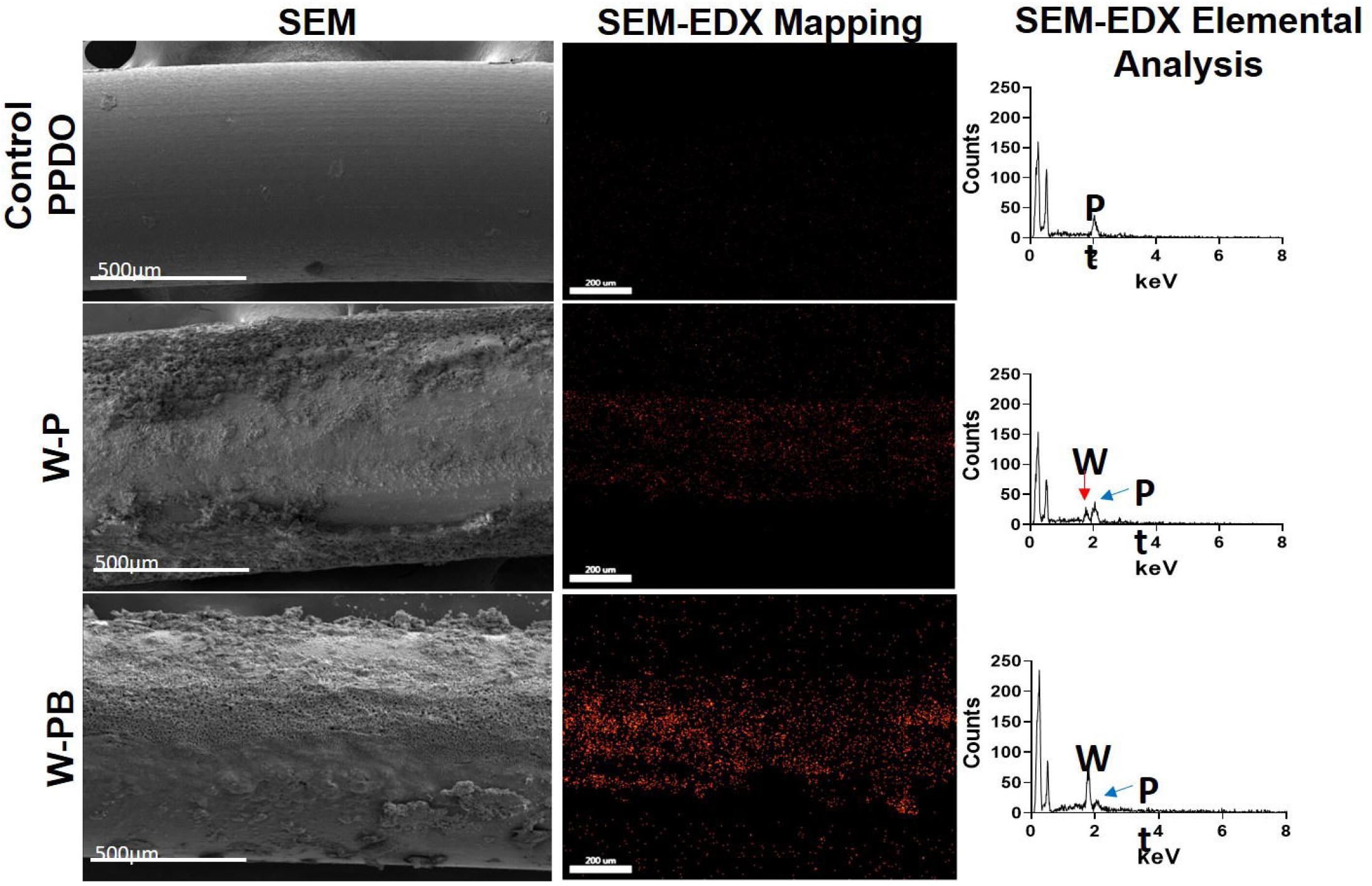
Scanning electron microscopy and tungsten mapping using energy-dispersive X-ray spectroscopy (EDX). Bare PPDO sutures had a smooth surface, and addition of polymers increased the roughness of the surface of the PPDO sutures, as shown by SEM. The peak at 1.77 keV confirms the presence of tungsten (W) for both W-P- and W-PB-coated sutures. A platinum (Pt) peak at 2.05 keV is also evident due to the sputter coat used for SEM.

**Fig 3.**
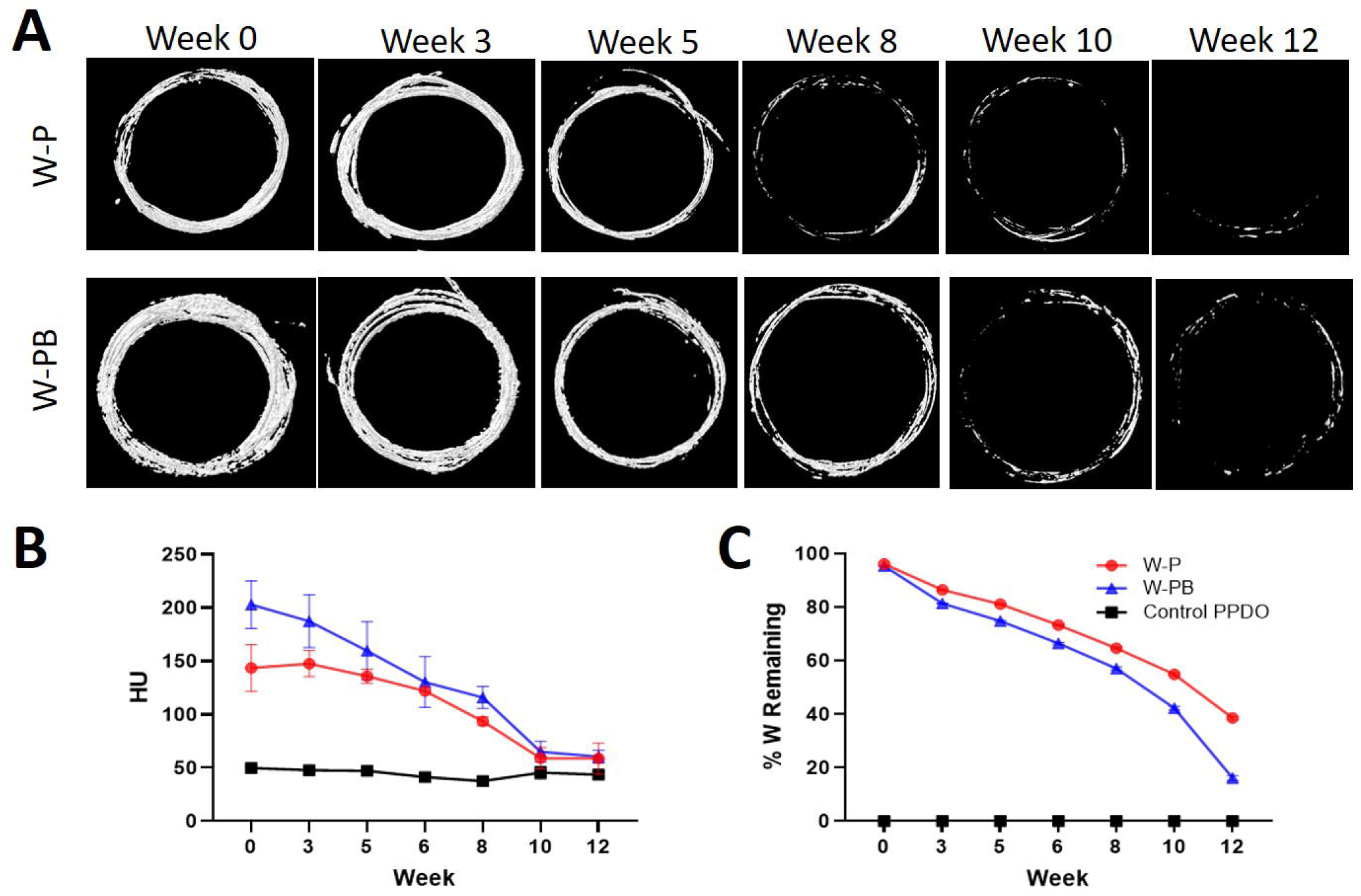
Longitudinal monitoring of the amount of tungsten and the radiopacity of W-P- and W-PB-coated sutures. (A) MicroCT images of sutures of W-P and W-PB from week 0 to 12. (B) Quantification of radiopacity in HU. (C) Percentage of W remaining over a span of 12 weeks as quantified in ICP-OES. Both W-P and W-PB showed sustained release of WNPs from the sutures; however, W-PB exhibited a drastic decrease of WNPs from week 8 to 12. This trend is consistent with the microCT images taken over the span of 12 weeks.

When processed using the same threshold levels, X-ray and microCT images showed that W-PB (2132.19 ± 323.44 HU) had higher radiopacity compared to W-P (99.78 ± 13.58 HU) and control PPDO (33.72 ± 8.15 HU), as shown in Fig 4. These results show that the PHB, PCL, and PVP polymer blend enhances the attachment of W on the surface of the PPDO sutures, thereby increasing the amount of WNPs loaded onto the PPDO sutures.

**Fig 4.**
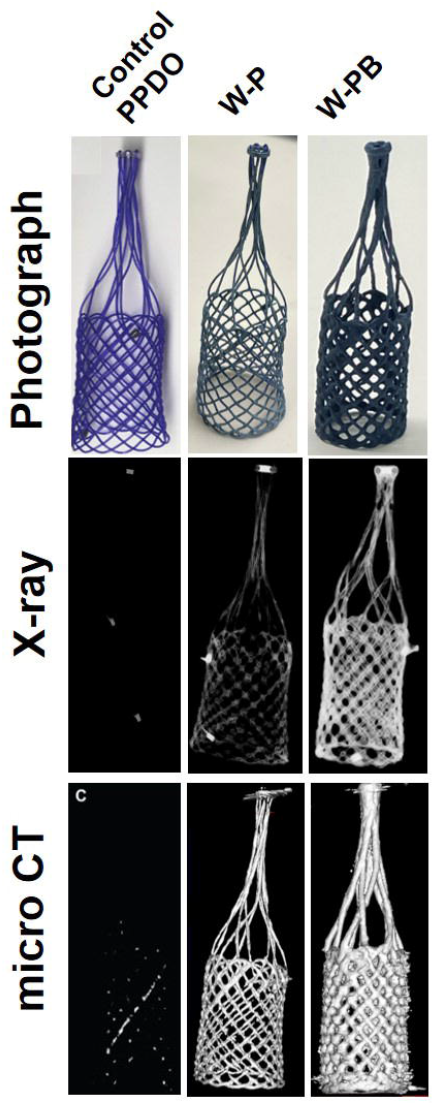
Comparison of the photograph, X-ray, and microCT images of control, W-P-coated, and W-PB-coated IVCFs. The additional polymers in W-PB-coated filters make them darker and thicker, with improved radiopacity on both X-ray and microCT compared to control PPDO and W-P-coated IVCFs.

Before proceeding with *in vivo* studies, we wanted to establish that the desired physicochemical and mechanical properties of PPDO sutures remain optimal upon the addition of W-P and W-PB. We compared the mechanical strength and melting temperatures of W-P- and W-PB-coated sutures to the control PPDO (Table 1). We found that the mechanical strength of the W-P (maximum stress, 37.04 ± 4.56 mPa; *p* = 0.5785) and W-PB (31.70 ± 4.12 mPa; *p* = 0.9773) sutures did not vary significantly from the control PPDO sutures (32.56 ± 5.63 mPa) at baseline. The melting temperatures of the sutures were also similar across treatments. These results are consistent with our previous studies of AuNP-coated^3^ and BiNP-coated^9^ IVCFs, in which addition of NPs did not alter the mechanical strength of the PPDO sutures.

### Image-guided deployment and long-term radiopacity of WNP IVCFs

The deployment of IVCFs in pigs was done under fluoroscopic guidance. Fluoroscopic images of the catheter and clot deployments of W-P- and W-PB-coated IVCFs are shown in Fig 5. W-PB filters showed improved visualization in *in vivo* fluoroscopic images as compared to W-P filters, which was expected given the increased WNP loading when the polymer blend was used. Clot deployments immediately followed filter deployment. WNPs and the PHB, PCL, and PVP polymers did not affect the expected clot-trapping efficacy of PPDO IVCFs; both W-P and W-PB filters were able to capture the administered clots.

**Fig 5.**
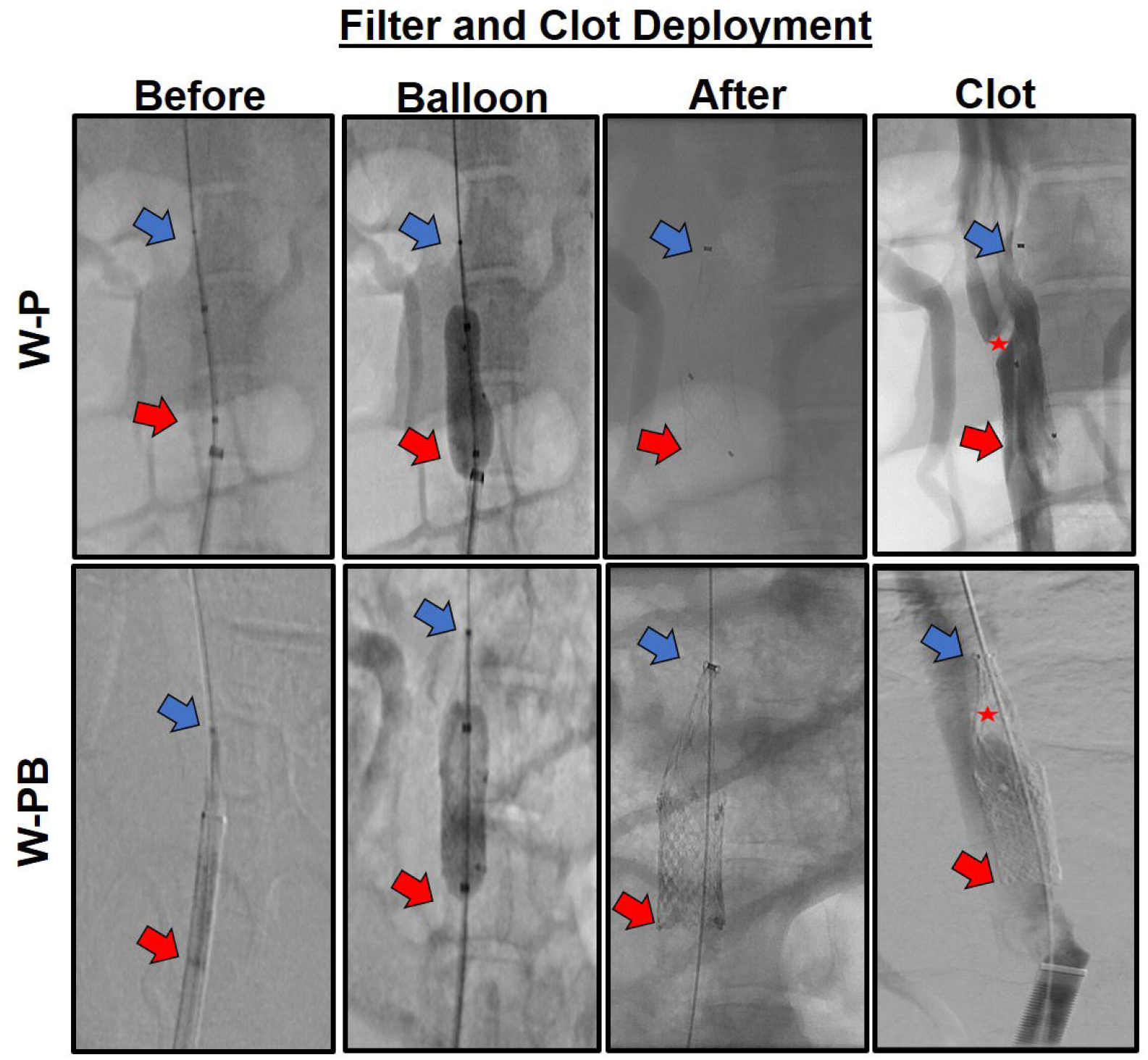
*In vivo* filter deployment and clot-trapping feasibility of W-P and W-PB filters. lntraoperative fluoroscopic imaging of catheter deployment (Before), IVCF deployment (Balloon), and implantation (After) are shown. Clot deployment (Clot) is demonstrated with contrast-enhanced imaging; successfully trapped clots are denoted with red stars. Blue and red arrows show the top and bottom of the filters, respectively.

Longitudinal CT monitoring showed that the W-PB filter had 2-fold higher HU intensity (838.64 ± 44.64 HU) compared to the W-P filter (404.49 ± 52.39 HU) at the time of deployment (Fig 6). Furthermore, initial *in vivo* CT monitoring showed 11-fold signal enhancement of W-PB (763.07 ± 330.9 HU) compared to the background (muscle, 69.7 ± 13.00 HU), while W-P was only 8-fold higher (469.80 ± 220.00 HU) than the background (muscle 57.22 ± 8.00 HU). Maximum HU values showed decreasing CT intensities throughout the 12 weeks for both W-P only and W-PB (Fig 6A). These results support that the additional polymer blends in W-PB allow for better WNP binding and more sustained release of NPs throughout the 12-week period compared to the W-P single-polymer blend.

**Fig 6.**
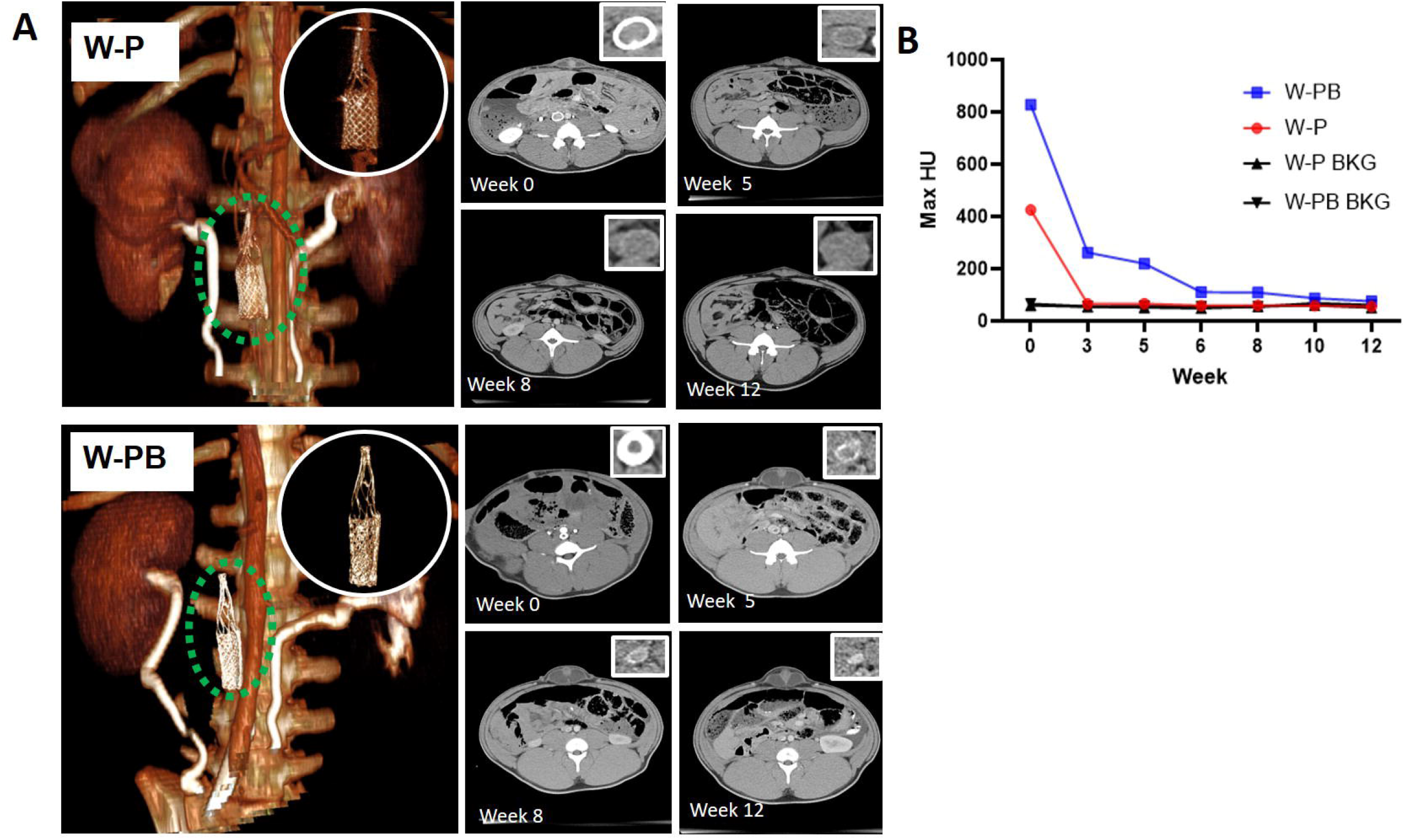
Longitudinal CT monitoring of IVCFs throughout the 12-week implantation period of W-P- and W-PB-coated IVCFs. (A) Coronal CT at the time of filter deployment (left) and axial images over time (right). (B) Maximum HU values of the IVCF ring compared to the background (BKG).

In our previous study using BiNP- and BiNP+PHB–coated IVCFs, we observed that the incorporation of the IVCFs in the vena cava wall resulted in neointima formation, which was most evident at weeks 5-6. In this study, we also attempted to measure the luminal area in response to the W-P- and W-PB-coated IVCFs. However, it was challenging to measure and account for the actual luminal area possibly due to beam-hardening artifact brought by the WNP coating. Moreover, the results we obtained from the longitudinal *in vitro* release experiments did not correspond to the results *in vivo*, where both W-P- and W-PB-coated IVCFs exhibited a drastic decrease in HUs as early as week 3 (Fig 6B). This may be due to the difference of environment *in vitro* versus *in vivo*. Although a stable release of WNPs was observed *in vitro* in PBS matrix, it is important to take into account the complexity of *in vivo* body fluids, oxidation, and enzymatic factors that may alter the performance of polymer-based materials.^22^

### *In vivo* safety assessment

Upon exsanguination of pigs, we discovered that the IVCF-affected areas for pigs implanted with W-PB-coated IVCFs were characteristic of a necrotic vessel (Fig 7). We then proceeded to harvest and quantify WNPs in blood and different organs. In humans, W is expected to translocate into the blood and circulate into the body after entering through ingestion or inhalation. Typically, W is rapidly excreted, but some may remain and be detected in the kidney, liver, spleen, and bone.^23^ For both W-P and W-PB pigs, we did not find traces of WNPs in the blood, heart, lungs, spleen, liver, kidneys, intestines, bladder, or stomach. However, we detected tungsten in W-P IVC upstream (0.149 ± 0.002 mg W / g tissue), IVC downstream (0.210 ± 0.003 mg W / g tissue), and IVC affected areas (0.175 ± 0.009 mg W / g tissue). An approximately 50-fold increase in W was detected in W-PB IVC upstream (5.273 ± 0.381 mg W / g tissue), IVC downstream (5.470 ± 0.706 mg W / g tissue), and IVC affected areas (6.151 ± 0.270 mg W / g tissue). Blood chemistry evaluation did not show any significant differences from the baseline values taken prior to filter deployment (Table S1), suggesting that the observed adverse effects were local to the affected IVC.

**Fig 7.**
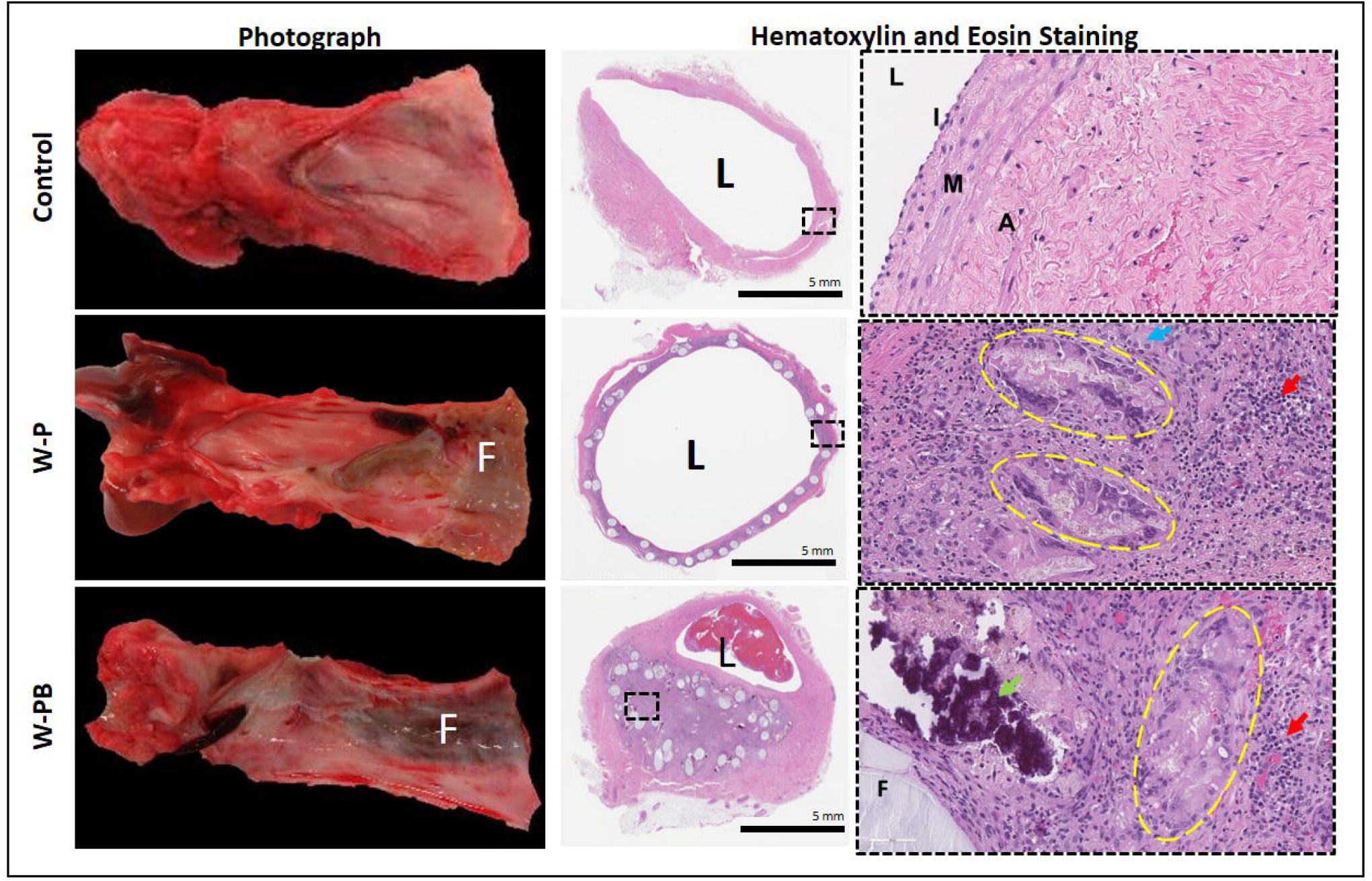
Gross necropsy and histology images of IVCs collected 12 weeks after implantation of W-P- and W-PB-coated IVCFs. At 40X, distinct inflammatory structures such as Langhans-type multinucleated cells (yellow dotted circle), epithelioid macrophages (blue arrow), lymphocytes (red arrows), and dystrophic mineralization (green arrow) can be seen surrounding the filter material. L: lumen, I: intima, M: tunica media, A: tunica adventitia, F: filter material.

Gross necropsy and histological examination of the IVC affected area showed incorporation of the filter into the walls of the IVC, which in turn resulted in the formation of a dense fibrous tissue and degradation of filter strands of the device (Fig 7). For both W-P- and W-PB-coated IVCFs, histological analysis (H&E) of the IVC affected tissue revealed fragments of NPs in the proximity of the incorporated filter strands. These filter fragments were visibly surrounded by macrophages and Langhans-type multinucleated giant cells, which are common features of granulomas and develop during an inflammatory response.^24^ Similar observations were seen for the W-PB affected area; however, the W-PB-coated filter evidently migrated and incorporated the exterior of the vascular lumen and into the tunica adventitia. Further analyses using immunofluorescence are recommended to visualize, characterize, and understand specific immune markers that are present in these tissues.

While current literature on the toxicity of WNPs is limited, studies on its toxicological effects have been inconsistent. For example, sodium tungstate nanoparticles were found to increase the frequency of apoptotic human-derived peripheral blood lymphocytes, alter cell cycle regulation, and reduce cytokine production at different doses and exposure times.^25^ Inhalation of tungsten oxide nanoparticles also induced cytotoxicity, morphological changes, and lung injury in golden Syrian hamsters through a pyroptotic cell death pathway.^26^ In contrast, tungsten oxide nanoparticles did not have any toxic effects on an *in vitro* 3D human airway epithelium model.^27^ One of the hypothesized toxic mechanisms of action of W substances is through the production of reactive oxygen species (ROS), which increase in concentration as the W particle size decreases.^28, 29^ While there is a lack of unifying data to explain the inflammatory response elicited from implantation of the W-P- and W-PB-coated IVCFs, further evaluation using immunohistochemistry may help us characterize and explain and possibly mitigate the inflammatory response associated with the necrotic tissues observed for both W-P- and W-PB-coated IVCFs.

## Conclusion

Tungsten NPs with layer-by-layer reinforcement of polymer (W-P) and polymer blends (W-PB) were successfully incorporated onto radiolucent, absorbable IVCFs, thereby enhancing radiopacity and overall clot-trapping efficacy. W-PB-coated IVCFs show improved radiopacity in fluoroscopy-guided deployment and long-term CT monitoring compared to W-P-coated IVCFs alone. The additional layers of PHB, PCL, and PVP in the W-PB coating improved WNP binding. However, this led to observed local cytotoxic effects in the IVC affected area of pigs implanted with W-PB-coated filters. As we have not observed any toxic effects from our previous NP-coated IVCFs, it is important to explore the different toxic mechanism underlying the inflammatory response observed with the implantation of W-P- and W-PB-coated IVCFs. A thorough toxicity assessment may help us mitigate the cytotoxic effects possibly brought about by the addition of both NP and polymer coatings before implantation of the IVCFs.

Our study on NP-coated absorbable IVCFs provides a foundation for developing nano-embedded medical devices. Experimentation with alternative materials and methods to develop radiopaque absorbable IVCFs is recommended. For example, 3D-printed IVCFs using NP-infused polymers may eliminate the need for layer-by-layer coating of sutures with or without the use of polymers, thereby also producing IVCFs with more uniform and sustained release *in vivo*.

## Author Contributions

Conceptualization – E.M.S.V., J.A.D., S.Y.H., M.P.M.; Data curation – E.M.S.V., J.A.D., M.B., K.A.C., G.M.C., G.C., M.C.J., N.F., M.P.M.; Formal analysis – E.M.S.V., J.A.D., M.B., K.A.C., B.G., G.M.C., G.C., M.C.J., N.F., M.P.M; Investigation – E.M.S.V., J.A.D., M.B., K.A.C., B.G., G.M.C., A.M., G.C., M.C.J., W.N., R.R.L., N.F., S.R.C., S.Y.H., M.P.M.; Supervision – M.P.M.; Project administration – J.A.D., M.P.M; Writing – original draft – E.M.S.V., M.P.M.; Writing – review and editing – E.M.S.V., J.A.D., M.B., K.A.C., B.G., G.M.C., A.M., G.C., M.C.J., W.N., R.R.L., N.F., S.R.C., S.Y.H., M.P.M.; Funding acquisition – S.Y.H., M.P.M. All authors have read and agreed to the published version of the manuscript.

## Conflicts of interest

There are no conflicts to declare.

## Acknowledgements

Funding for this work was provided by the National Institutes of Health (NIH)/National Heart, Lung, and Blood Institute (5R01HL141831-05 and 1R01HL159960-01A1; to M.P.M.); the NIH/National Cancer Institute (P30CA016672; used the Research Animal Support Facility and Small Animal Imaging Facility); and the Dunn Foundation. We would like to thank Adient Medical, headed by Dr. Mitch Eggers and supported by Stephen Dria, for providing the uncoated PPDO IVCFs. Dr. James Gu from the Houston Methodist Research Institute Electron Microscopy Core helped us obtain SEM/EDX data. TEM imaging was done by Kenneth Dunner at the MD Anderson High Resolution Electron Microscopy Facility. We are also grateful for the support of the large animal veterinary and the Small Animal Imaging Facility staff. Lastly, we would like to thank Ms. Sunita Patterson from the MD Anderson Cancer Center Research Medical Library Editing Services for editing our manuscript.

## Notes

### Competing Interest Statement

The authors have declared no competing interest.

