## Supplementary Table 1 for "Image-guided deployment and monitoring of a novel tungsten nanoparticle–infused radiopaque absorbable inferior vena cava filter in pigs"

Table S1. Serial hematologic parameters of pigs implanted with (A) W-P IVCF and (B) W-PB IVCF.

| A | Sodium (mEq/L) | Potassium  (mEq/L) | Chloride (mEq/L) | Creatinine (mg/dL) | WBC  (10^3^/µL) | Hemoglobin (g/dL) | Platelets (10^3^/µL) | PT  (s) | PTT  (s) | AST (U/L) | ALT  (U/L) | pH level | pCO_2_  (mmHg) | SO_2_  (%) |
| --- | --- | --- | --- | --- | --- | --- | --- | --- | --- | --- | --- | --- | --- | --- |
| Baseline | 131.3 | 4.8 | 96.2 | 1.15 | 13.85 | 9.4 | 480 | 12.8 | 38.1 | 16 | 24 | 7.454 | 48.6 | 100 |
| Week 3 | 138.1 | 4.02 | 103.8 | 1.38 | 9.61 | 9.7 | 451 | 12.9 | 37.7 | 19 | 33 | 7.446 | 54.6 | <> |
| Week 5 | 137.8 | 4.13 | 104.4 | 1.46 | 9.34 | 9.4 | 446 | 13 | 39.7 | 17 | 35 | 7.393 | 51.8 | <> |
| Week 6 | 138.7 | 4.07 | 104.2 | 1.64 | 8.75 | 9.5 | 390 | 13.1 | 37.8 | 20 | 36 | 7.396 | 53.7 | <> |
| Week 8 | 137.4 | 3.96 | 102.1 | 1.69 | 9.37 | 9.1 | 389 | 12.9 | 40.7 | 21 | 39 | 7.439 | 51.4 | 99% |
| Week 10 | 141.8 | 3.9 | 104.3 | 1.86 | 8.95 | 9.1 | 344 | 12.8 | 37.7 | 18 | 38 | 7.458 | 52 | <> |
| Week 12 | 142.3 | 4.2 | 106.8 | 1.95 | 9.03 | 9.6 | 342 | 12.4 | 38.3 | 24 | 37 | 7.402 | 55.6 | 100 |

| B | Sodium (mEq/L) | Potassium  (mEq/L) | Chloride (mEq/L) | Creatinine (mg/dL) | WBC  (10^3^/µL) | | Hemoglobin (g/dL) | Platelets (10^3^/µL) | PT  (s) | PTT  (s) | AST (U/L) | ALT  (U/L) | pH level | pCO_2_  (mmHg) | SO_2_  (%) |
| --- | --- | --- | --- | --- | --- | --- | --- | --- | --- | --- | --- | --- | --- | --- | --- |
| Baseline | 140.2 | 4.72 | 99.4 | 1.18 | | 17.82 | 10.3 | 545 | 12.4 | 39.4 | 23 | 30 | 7.457 | 52.8 | 100 |
| Week 3 | 140.3 | 3.98 | 100.5 | 1.52 | | 15.57 | 9.3 | 670 | 12.1 | 36.4 | 19 | 27 | 7.485 | 52.6 | 100 |
| Week 5 | 142.1 | 3.95 | 102.7 | 1.82 | | 16.33 | 9.8 | 579 | 12.2 | 33.3 | 18 | 29 | 7.477 | 47.2 | 100 |
| Week 6 | 138.9 | 3.96 | 100 | 1.67 | | 14.24 | 9.1 | 632 | 12.3 | 40.2 | 17 | 22 | 7.493 | 48.9 | 100 |
| Week 8 | 144.7 | 3.9 | 106.1 | 1.75 | | 12.17 | 9.8 | 487 | 12.3 | 33.3 | 16 | 31 | 7.5 | 49.3 | <> |
| Week 10 | 143.4 | 3.79 | 102.1 | 1.82 | | 9.88 | 9.5 | 465 | 12.4 | 37 | 17 | 32 | 7.525 | 47.6 | <> |
| Week 12 | 142.1 | 4.06 | 102.7 | 2.03 | | 11.26 | 9.5 | 524 | 12.5 | 37.4 | 19 | 28 | 7.497 | 48.1 | <> |
